# DIRECT CALCULATION OF VARIABLE-RESOLUTION MAPS BY FULLY ANALYTIC APPROXIMATIONS TO ATOMIC IMAGES

**DOI:** 10.1101/2025.07.18.664315

**Authors:** Alexandre G. Urzhumtsev, Vladimir Y. Lunin

## Abstract

Crystallographic maps are typically computed as Fourier series using complex-valued structure factors, with a single set of coefficients applied across the entire region, and associating a single resolution value with the whole map. In cryo-electron microscopy, however, experimental maps often have variable local resolution. As a consequence, applying the same procedure to compute model maps for comparison with the respective experimental ones becomes complicate. An alternative strategy involves representing model maps as sums of atomic images. They are oscillatory functions of the distance from the atomic center and depend on atomic displacement parameters and local resolution. To calculate such maps, we propose a stand-alone program which uses the shell-function approximation of atomic images making them analytic functions of all its parameters including the local resolution associated to each atom.

**Synopsis:** Crystallographic and cryo electron microscopy maps with variable resolution can be calculated as a sum of analytic shell-function approximations to atomic images using the developed stand-alone python script.

## 1. Introduction

Crystallographic maps are typically computed as Fourier series using complex-valued structure factors, with a single set of coefficients applied across the entire map. Consequently, a uniform resolution is usually assigned to the map, although directional anisotropy in resolution can be analyzed (*e.g*., Urzhumtseva *et al*., 2013; Tickle *et al*., 2016, and references therein). An alternative approach involves computing the map as a sum of atomic contributions at limited resolution. While this procedure complicates the inclusion of non-atomic components such as bulk solvent, it offers several advantages. Notably, it is particularly well-suited for generating maps of electrostatic scattering potential in cryo electron microscopy (cryo-EM), where resolution often varies across different regions (*e.g*., Cardone *et al*., 2013), leading to the concept of local resolution (Kucukelbir *et al*., 2014; Vilas *et al*., 2018, and others).

These atomic contributions are atomic images computed at a given resolution *d*, with corresponding atomic displacement parameter (ADP) and occupancy values. Therefore, to reproduce the experimental map as a sum of such images, each atom of the model must be assigned an additional parameter, namely the local resolution of the map in its vicinity (Urzhumtsev & Lunin, 2022a).

Atomic images used to calculate a map are oscillating functions of the distance from the atomic center. These images, or rather approximations to them, can be computed either numerically or as analytic functions of ADP at a given resolution (*e.g*., Diamond, 1971, Lunin & Urzhumtsev, 1984; Chapman, 1995; Mooij *et al*., 2006; Chapman *et al*., 2013; DiMaio *et al*., 2015; Sorzano *et al.*, 2015; Pintilie *et al*., 2020; and references therein). When atomic scattering factors are approximated by a sum of Gaussian functions (*e.g*., Doyle & Turner, 1968; Agarwal, 1978; Waasmaier & Kiefel, 1995; Peng, 1999; Grosse-Kunstleve *et al*., 2004; Brown *et al*., 2006), the corresponding atomic images can be written as analytic functions of both ADP and resolution (Lunin & Urzhumtsev, 2022a). This can be done representing them as series of especially developed spherically symmetric three-dimensional functions, *Ω*_3_(**r**; *µ, v*), which depend on two scalar parameters, μ and *v* (Urzhumtsev & Lunin, 2024), defining the radius and the width of the spherical shell within which the function values are concentrated.

This manuscript presents a stand-alone program that calculates maps with potentially variable resolution across the regions as a sum of analytic approximations to atomic images.

## 2. Methods and algorithms

### 2.1 Shell functions

The approximation of atomic scattering factors, and thus atomic densities, by Gaussian functions is important because it renders them analytic functions of ADP. This property arises from the invariance of the class of Gaussian function under convolution with other Gaussians. However, atomic images at limited resolution, which are inherently oscillatory in three-dimensional space, cannot be approximated by Gaussian functions. Instead, they can be represented by shell functions *Ω*_3_(**r**; *µ, v*), a class of spherically symmetric functions whose values are concentrated within a shell of prescribed width *v* centered at distance μfrom the origin. This class is also invariant under convolution with Gaussians, preserving analytic dependence on ADP (Urzhumtsev & Lunin, 2024). Gaussian functions represent a particular case of *Ω*_3_(**r**; *µ, v*) with μ= 0.

By approximating the interference function in three-dimensional space with shell functions and the atomic scattering factors with Gaussians, atomic images can be expressed as analytic functions of all atomic parameters, including the local resolution *d* associated with each atom (Urzhumtsev & Lunin, 2022a). While this approach is computationally more demanding than directly using shell approximations to atomic images precomputed at a fixed resolution, it enables a more detailed analysis of the model-to-map correspondence. It also opens the possibility for analytic refinement of local resolution values, now associated with atoms. Although potentially less practical for fast refinement procedures with fixed resolution, this method offers substantial benefits for advanced model assessment and resolution mapping.

### 2.2 Procedure and its parameters

To implement the procedure described above, a stand-alone program, *ModQMap*, has been developed. The program takes as input an atomic model and a file containing the respective atomic scattering factors approximated by Gaussian functions. Based on this information, it generates a map of the corresponding distribution, such as electron density or electrostatic scattering potential.

Each atom, in addition to its coordinates in the crystal (for crystallography) or pseudo-crystal (for cryo-EM), occupancy, and ADP, is also assigned an individual value of local resolution, *d*. This local resolution can be stored directly in the atomic records. Alternatively, it may be assigned as a global value shared by all atoms, which is convenient for crystallographic applications, or extracted from a map of local resolution (*e.g*., Dai *et al*., 2023). In the latter case, the local resolution at the nearest grid point is assigned to each atom.

The procedure employs the *Ω*_3_(**r**; *µ, v*) approximation to the three-dimensional normalized interference function π_3_(**r**) as described by Urzhumtsev & Lunin (2024). Using this approximation, each atomic image is computed within a spherical region of radius *R*_max_ centered at the atomic position. Increasing *R*_max_ improves the map accuracy but increases the computation time. Previous analyses (Urzhumtsev *et al*., 2023) have shown that a better accuracy can be achieved choosing *R*_max_ as integer or half-integer multiple the resolution *d* associated with the given atom, *R*_max_ = *kd*. For sufficiently accurate maps, the factor *k* equal to 2.0 or 2.5 is typically sufficient (Urzhumtsev & Lunin, 2022b). Since each *Ω*_3_(**r**; *µ, v*) term contributes within a spherical shell of finite width, a few additional terms with *μ* > *R*_max_ should be included to ensure map accuracy.

The grid parameters used for map calculation, namely, the map boundaries and grid step, can either be specified directly in the input parameters or taken from a reference (experimental) map, if provided.

Although cryo-EM calculations are typically carried out within orthogonal pseudo-cells, the procedure is general and supports maps and models defined in arbitrary unit cells. However, the current version of the program does not apply symmetry operations. Therefore, all atoms contributing to the computed portion of the map must be explicitly included in the input atomic model.

The calculated map can be computed in absolute or in sigma-valued scales, or be optimally scaled to match the experimental map either globally or within the molecular region defined as the set of grid points where the model contributes.

If an experimental map is provided, the program automatically compares the calculated and experimental maps, both over the entire volume and within the molecular region.

Additionally, if a map-like file of local resolution is supplied, the program can generate an output model file in which each atom is annotated with its corresponding local resolution value. This facilitates downstream analysis and further computations.

## 3. Software

### 3.1 System and software requirements

The program is stand-alone and does not require installation. It is written in basic *Python*3 and relies on the standard *NumPy* library. The program, along with associated data files, is freely available upon request from one of the authors (AGU).

### 3.2 Input data

The program requires a text file, *ModQMap*.*dat*, containing input parameters that specify both the input / output files, and the calculation settings. All parameters are defined by keywords, and default values are available for most of them. An example input file is provided, including the full list of parameters and explanatory comments on their usage, and possible and default values.

The atomic model file can be provided in either *pdb* or *cif* format. Atomic records can be completed by the values of the associated local resolution. If these values are available for individual atoms, they can be included directly in the model file. For *cif* format, currently they should be specified using the keyword *_atom_site.resolution* ; for *pdb* format, the resolution values should occupy columns 67–72 of atomic records, otherwise free except special cases. Unit cell parameters may be defined, or redefined via *ModQMap*.*dat*.

Two more input files with standard information are required. These files are maintained externally from the program, making it straightforward to update or replace them as needed without modifying the code. The first file contains coefficients of Gaussian approximations to the atomic scattering factors. Several versions of this file are provided, and the desired one can be selected via input parameters. The computation time is approximately proportional to the number of Gaussians used in the approximation. However, while simpler approximations may reduce computation time, they may compromise map accuracy.

The second file contains the coefficients for the *Ω*_3_(**r**; *µ, v*) approximation of the three-dimensional interference function as given by Urzhumtsev & Lunin (2024). The current approximation to the interference function enables calculation of atomic images up to a distance of *R*_max_ = 20*d* where *d* is the local resolution.

Optionally, the user can provide an experimental map and a local resolution map as additional inputs, both in either in *mrc* or in *cns* formats, enabling further comparison and analysis.

### 3.3 Output data

The calculated map can be saved either in *mrc* or in *cns* formats.

The program creates a *log*-file which mirrors input information and provides with various statistics.

If requested by the input parameters, the atomic model completed with the associated local resolution values can be output in the same format as the input model, either *pdb* or *cif*.

## 4. Practical application to macromolecular studies

The structure of the nucleosome-sirtuin complex (PDB entry code 8of4) is composed of 9 proteins and 2 nucleic acids containing in total more than 14,000 non-hydrogen atoms (Smirnova *et al*., 2023). The composite map of the electrostatic scattering potential (emd_16845.map) shows a significant resolution heterogeneity (Fig. 1a). The local resolution of the map has been estimated using *CryoRes* (Dai *et al.*, 2023). Fig. 2 shows the distribution of the local resolution values associated with atoms; it varies near continuously between 4 and 5 Å for most of atoms. For about 300 atoms, the local resolution extracted from the resolution map was 100 Å as formally assigned by *CryoRes* for too blurred map regions.

**Fig. 1.**
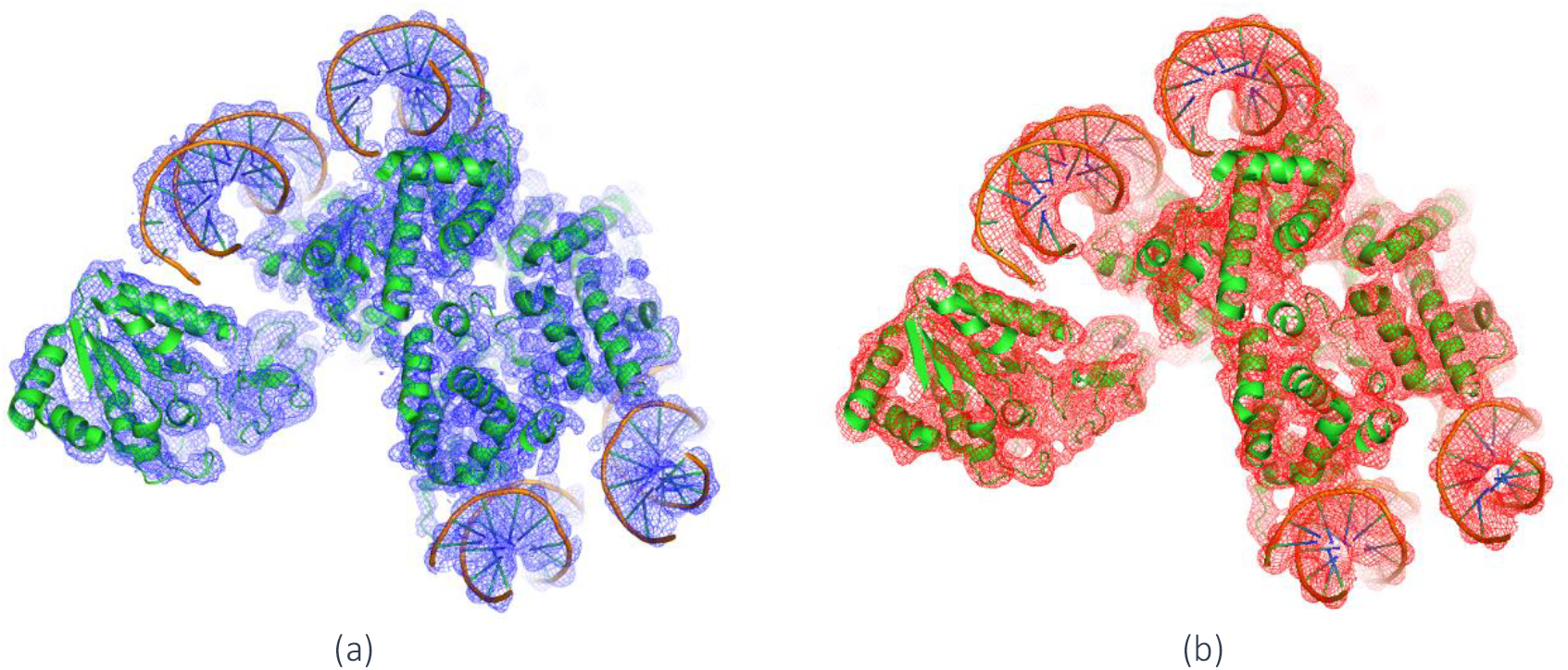
Experimental cryo-EM (a) and calculated (b) variable-resolution maps of the nucleosome-sirtuin complex. Figures are prepared with *Pymol* (Schrödinger & De Lano, 2020).

**Fig. 2.**
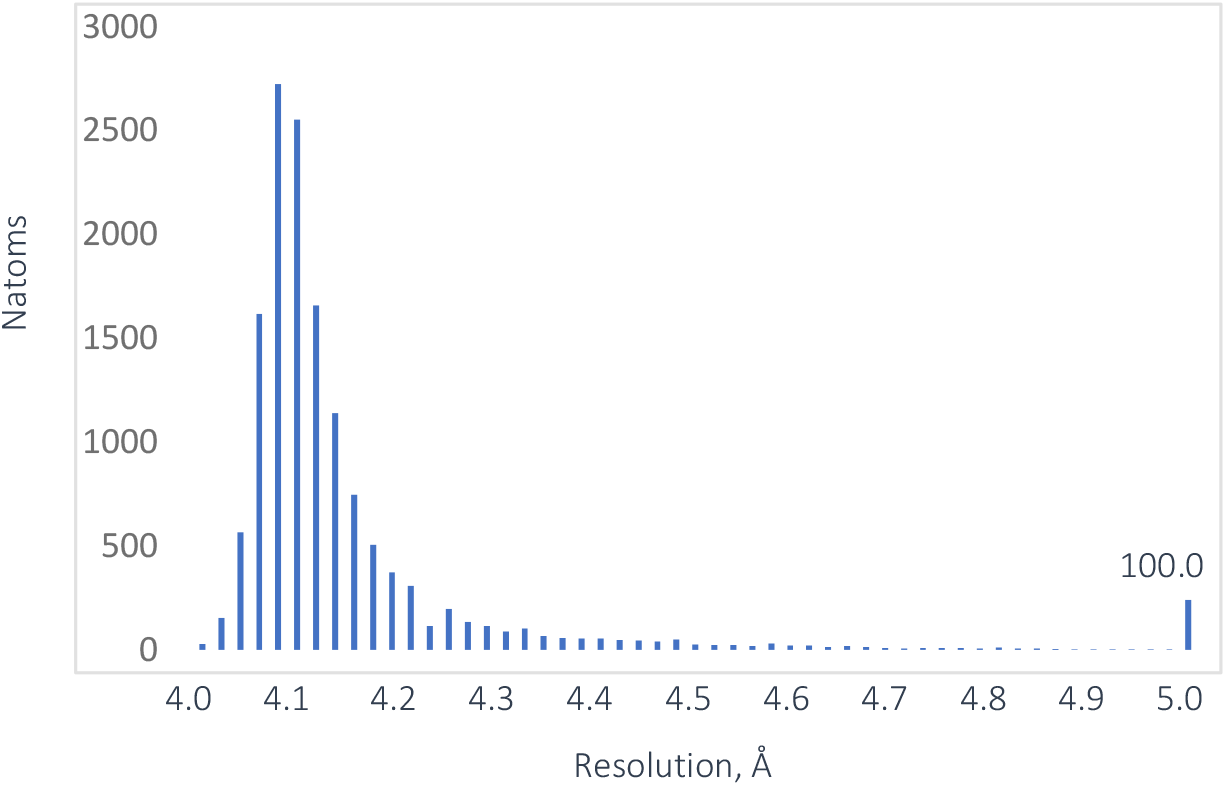
Distribution of the local resolution values associated with the atoms of the model of the nucleosome-sirtuin complex

To reproduce the experimental map, we used *ModQMap* which is especially adapted to such a situation when atomic images significantly vary according to their displacement parameters and resolution values. First, we tried a fast-and-dirty calculation with *R*_max_ = *d* and substituting 100 Å by 7 Å, already sufficient to drastically blur atomic images. This calculation took only about a minute. Then we recalculated the map increasing significantly the parameters values, atomic radius up to *R*_max_ = 2.5*d*, required for high-accuracy reproduction of images of individual atoms, and the resolution truncation value from 7 to 10 Å. This practically does not change overall statistical values indicating that, when a high accuracy is not required, including only a couple of ripples into calculation of atomic images is sufficient while still important (Urzhumtsev & Lunin, 2022a). Fig 1b shows the calculated map.

## 5. Discussion

Structural macromolecular studies require various computer tools adapted to different goals and answering different questions. While modern workflows rely on large software packages, stand-alone programs retain important advantages, most notably, offering users greater transparency into the underlying calculations and flexibility to tailor the software to specific needs. Naturally, such stand-alone tools are expected to remain compatible in file formats with major packages.

We present here an example of such a tool and one of its applications to macromolecular structure analysis, namely, the cryo-EM studies of the nucleosome-sirtuin complex. Developed in *Python* rather than compiled languages like *Fortran* or *C*++, the program is not optimized for speed and is therefore not intended for integration into iterative refinement procedures. Instead, as the example shows, with its extensive parameterization, it is particularly well-suited for methodological investigations and targeted analyses of experimental images in macromolecular crystallography and in cryo electron microscopy.

## Acknowledgments

The authors thank G. Papai for help with the nucleosome data. AU acknowledges the French Infrastructure for Integrated Structural Biology FRISBI [ANR-10-INBS-05] and the support and the use of resources of Instruct-ERIC through the R&D pilot scheme APPID 2683. VL acknowledges support of KIAM RAS FFMN-2025-0024.

